# The dynamic relationship between pupil dilation and neural surprise in natural language comprehension

**DOI:** 10.64898/2026.06.12.731866

**Authors:** Quirin Gehmacher, Aaron Kaltenmaier, Juliane Schubert, Nathan Weisz, Clare Press

## Abstract

Predictive processing accounts hold that the brain continuously generates predictions and updates internal models from error signals, but it remains unclear whether prediction representation and model-updating reflect a single graded computation or computationally distinct operations implemented in dissociable systems. We recorded magnetoencephalography (MEG) and pupillometry while participants listened to continuous natural speech, quantifying word-by-word lexical surprise and semantic prediction error with a GPT-2 model. Broadly speaking, lexical surprise was tracked in a predominantly graded fashion, whereas the response to semantic error was better captured by a rectified linear (ReLU) gating function - i.e., that responds only when the error exceeds a recent contextual baseline. There were also some interesting dissociations in the strength of these relative effects. The auditory cortex tracked lexical surprise in a graded, continuous manner, with later cortical stages reflecting semantic error. By contrast, tonic pupil diameter and source-localised brainstem responses were predominantly captured by a gated response to semantic error. A pupil-coupling analysis confirmed that this gating signature was statistically shared between pupil-linked arousal and brainstem-localised, but not cortical, responses. Together, these findings reveal a division of labour in which the cortex maintains a continuous, high-fidelity map of predictive information while a pupil-linked model-update system acts as a selective gate, engaged specifically by meaning-disrupting events that have crossed a threshold level of error. This asymmetry suggests that signals serving continuous parsing of prediction error and signals serving model revision may be computationally dissociated, with a range of implications for our understanding of the intrinsic interactions between mechanisms serving perception and learning.

## Introduction

Predictive processing accounts posit that learning about the structure of the world shapes our processing of perceptual input in a way that optimises future learning ^1–4^. Error signals between predictions and incoming sensory evidence are proposed to be computed even at the earliest stages of processing ^5–7^, and to, in turn, recruit brainstem neuromodulatory systems - including the locus coeruleus and basal forebrain - to optimise model-updating ^8–10^. The nature of this iterative loop between perception and learning remains poorly understood in humans, however, in part because quantifying precise levels of surprise in naturalistic input has been difficult, and because the subcortical systems implicated in model-updating are difficult to access non-invasively.

Advances in large language models (LLMs) have substantially reduced the first of these difficulties. Trained to predict upcoming words from context, LLMs can be used to quantify prediction error in continuous narrative input through contextual word-level estimates of lexical surprise - the (un)likelihood of a specific word given its context - and semantic prediction error - the mismatch between a word’s meaning and the meaning expected from the preceding context. Cortical responses have been shown to track these signals during natural language comprehension across a distributed language network ^11–13^. Canonical predictive-processing frameworks are often cast as if organisms continuously and iteratively use all error signals to update internal models. A recent proposal, however, holds that the perception-learning interface may operate qualitatively differently depending on the magnitude of prediction error ^14^: only errors that exceed a context-sensitive threshold should trigger neurochemical processes that optimize model updating. This account implies a functional dissociation between the sensory representation of prediction error and the engagement of update mechanisms - with early cortical systems tracking contextual prediction error in a graded manner, while brainstem-linked neuromodulatory systems operate according to a rectified, gating-like computation. Such gated computations would subsequently be expected to influence processing throughout the hierarchy.

Probing this dissociation non-invasively is also complicated by the difficulty of resolving deep neural sources with time-resolved imaging. MEG has recently been purported to reveal non-cortical structures during speech processing - for instance, recovering cerebellar responses to rhythmic speech ^15^ - yet any single deep-source reconstruction remains hard to interpret in isolation. We therefore reasoned that a claim about brainstem-linked gating would be strengthened by a convergence between independent measures. The brainstem is favourably placed in this respect: pupil dilation provides an indirect peripheral readout of brainstem-linked neuromodulation ^16,17^, offering a handle on the same system that is independent of any source reconstruction. Pupillary measures reflect many processes as well as brainstem activity, and source reconstruction of deep structures is noisy, but convergence between such a peripheral index and source-localised MEG would speak to brainstem-linked computation with a confidence that neither could carry alone. Yet the peripheral side of this argument also rests on an untested assumption: it is not known whether pupil dilation tracks LLM-derived contextual surprise during natural speech in the first place. If it does, the convergence logic predicts a specific dissociation - that a thresholded function should explain processing in pupil-linked and brainstem signals, but early cortical representations of error should be more graded. Furthermore, it is unknown how these functions might relatively explain lexical and semantic forms of surprise. One possibility of interest to us was that gating functions playing a role in model-updating might preferentially fit semantic errors, which are arguably most behaviourally-relevant in natural human communication.

To answer how these structures interact in shaping the perception-learning interface, we recorded magnetoencephalography (MEG) and pupillometry concurrently while participants listened to narrative speech, source-localising the neural signal across the cortex as well as to the brainstem. We used temporal response functions (TRFs) to estimate how contextual surprise, derived from GPT-2 (following ^12^), was represented across these signals, comparing a conventional graded model with a gated one in which surprise was half-wave rectified relative to its recent history. This gated analysis was inspired by rectified linear unit (ReLU) activations in artificial neural networks as a simple formalisation of thresholded model-updating. A mediation analysis then tested whether pupil-linked arousal was coupled to the brainstem-localised rather than the cortical surprise response. Across most analyses, we replicated a basic pattern whereby lexical surprise was better modelled by a graded than gated response, yet semantic error showed the better fit to a gated signature. As well as these similarities, there were interesting differences in the relative strength of these effects across the neural and pupillometry measures. The interaction pattern in early sensory processing was predominantly driven by tracking of lexical surprise in a graded rather than thresholded form, with little tracking of semantic error. Later cortical regions - e.g., parietal and frontal structures - were more sensitive to semantic error modelled in both manners. In contrast, there was a threshold-sensitive profile to semantic prediction error in source-localised brainstem responses that was mirrored in pupil dilation, with a mediation analysis establishing that pupil-size was coupled to the brainstem-localised response to gated semantic prediction error. The pupillary measure was not coupled to cortical responses. Later cortical profiles become more gated, likely related to downstream influences of these signals reflected in the pupillary response and brainstem-localised analyses. Together, these findings indicate a division of labour in which the cortex continuously represents lexical predictive information, whereas a pupil-linked update system acts on it selectively, engaged when meaning-disrupting events signal the need to update one’s model of the world.

## Results

Twenty-eight participants listened to continuous German-language audiobook segments while MEG and pupillometry were recorded concurrently (Figure 1A-B). The pupil time series was decomposed into the raw signal and its first temporal derivative, which index dissociable components of pupil-linked neuromodulation ^16,17^. We submitted audiobook transcripts to a German GPT-2 model to obtain, for each word, a context-dependent prediction of the upcoming word (Figure 1C). From this information, we derived two contextual prediction-error measures: lexical surprise and semantic prediction error (Figure 1D; following ^12^).

**Figure 1:**
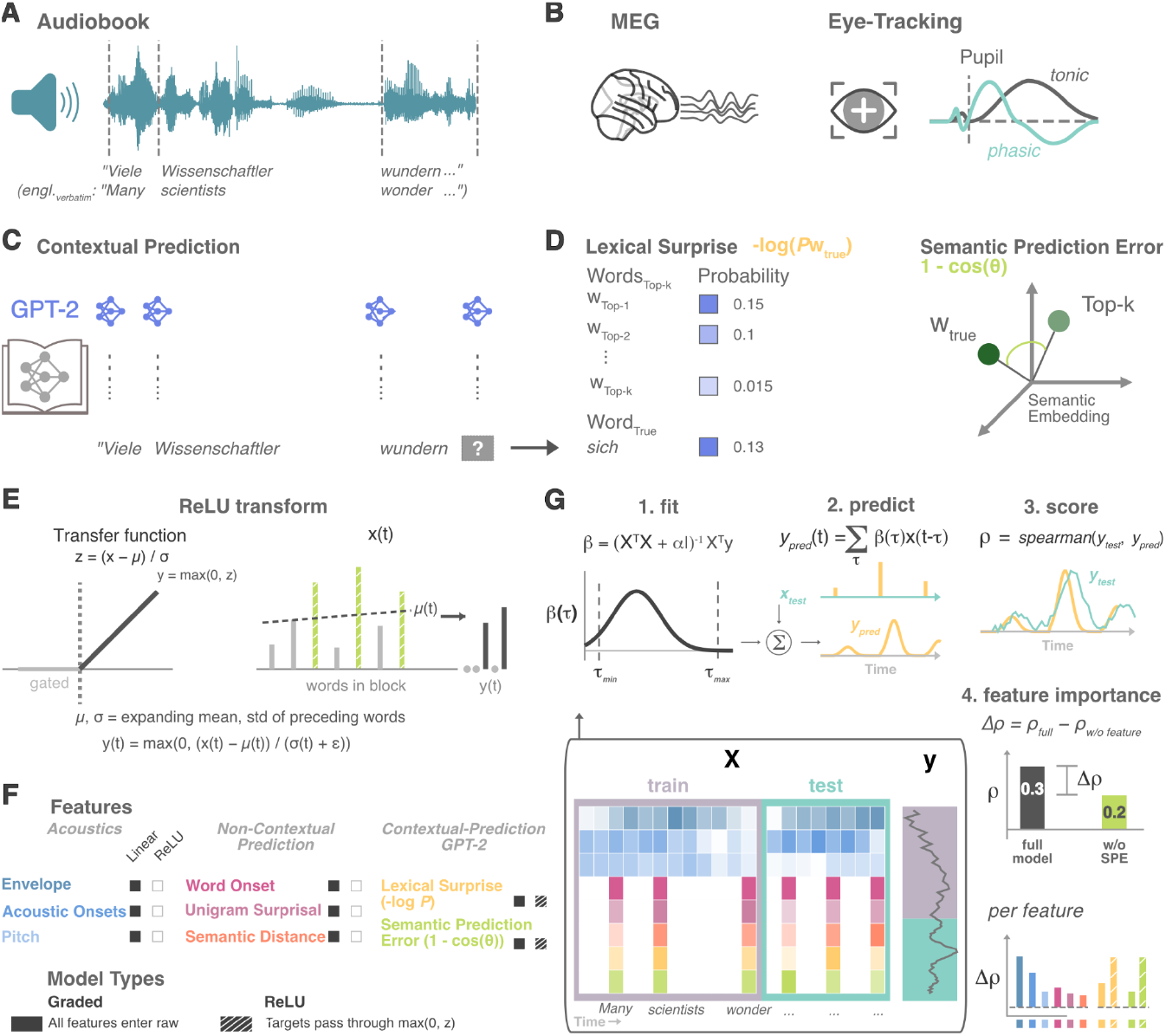
Experimental and analytical framework. (A) Participants listened to continuous German-language audiobook segments while word timings were aligned to the speech waveform; English translations are shown for illustration. (B) MEG and Eye-tracking were recorded concurrently, and pupil time series were decomposed into the raw pupil signal and its first temporal derivative as putative indices of tonic and phasic pupil-linked activity. (C) Transcript words were submitted to a German GPT-2 model to generate context-dependent predictions about upcoming words. (D) From these predictions, we derived two contextual prediction-error measures at the word level: lexical surprise, defined as the negative log probability of the observed word, and semantic prediction error, defined as the cosine distance between the embedding of the observed word and a probability-weighted average of likely next-word embeddings. (E) To test whether update-related responses are gated rather than graded, we additionally transformed predictors using a history-dependent rectified linear unit (ReLU), which standardized each word’s surprise relative to the recent local distribution and retained only positive deviations above that contextual baseline. Specifically, it returns as 0 all values below that baseline and only takes on a value above it. (F) Encoding models included acoustic features, non-contextual lexical features, and GPT-2-derived contextual prediction features, entered either in graded form or, for selected predictors, in ReLU-transformed form. (G) Temporal response functions were fit on training data and evaluated on held-out data by correlating predicted and observed responses; feature importance was quantified as the loss in predictive performance after removing each feature from the full model.

To test whether update-related responses are gated rather than graded, contextual prediction regressors were additionally entered after a history-dependent rectified-linear (ReLU) transform, retaining only positive deviations above a recent local baseline (Figure 1E). Prediction signals were then estimated using temporal response functions, which model each recorded signal as a time-resolved linear combination of predictors aligned to word onset (Figure 1F-G). Acoustic features (envelope, acoustic onsets, pitch) and non-contextual prediction regressors (word onset, unigram surprisal, semantic distance) were included in every model, so that any contribution of contextual prediction is estimated over and above lower-level structure. For each predictor, a full model fit on three of the four blocks was used to predict the signal in the held-out block, and prediction accuracy was quantified as the Spearman correlation between the predicted and observed time courses. A predictor’s contribution was then quantified as the drop in this accuracy when that predictor was removed from the full model (Figure 1G).

### Auditory cortex encodes lexical predictions in a graded, rather than gated, manner, with semantic error reflected in higher order structures

To sum, the cortex, as a whole, predominantly tracked lexical surprise in a graded form, with the graded formulation outperforming the gated one. It also tracked semantic prediction error, with no significant difference between the graded and gated formulations. When separated by region, we found that the auditory cortex tracked most clearly graded lexical surprise, with both graded and gated semantic error regressors increasingly contributing to prediction accuracy as moving to higher order structures, such as the frontal cortex.

We quantified each predictor’s contribution as the drop in prediction accuracy when it was ablated from the full model, which included acoustic, non-contextual, and the contextual prediction regressors (lexical surprise, semantic prediction error). Spatial inference used threshold-free cluster enhancement across the cortical parcellation (TFCE; Smith & Nichols, 2009), with significance assessed against a permutation null at α = .05 corrected. Reported peak parcels and cluster extents are descriptive - they indicate where effects were most strongly expressed rather than implying parcel-specific significance (see Methods).

When looking across the cortex as a whole, both contextual prediction regressors (i.e. lexical surprise and semantic prediction error) improved cortical encoding over the acoustic and non-contextual prediction regressors. Lexical surprise was significant across a broadly distributed, bilateral peri-Sylvian network (207/360 parcels; smallest *p* = .0001), and semantic prediction error over a smaller, predominantly left fronto-temporal extent (30/360 parcels; smallest *p* = .015), conceptually replicating ^12^. The gated (ReLU) formulation did not improve fit for either predictor. For lexical surprise the graded model fit better, over a right-lateralised motor and opercular extent (86/360 parcels, *p* = .001); for semantic prediction error the gated-graded contrast showed no significant effect (0/360 parcels). The lower-level nuisance features improved encoding as expected (envelope, acoustic onsets, and unigram surprisal; Figure 2A-B).

**Figure 2.**
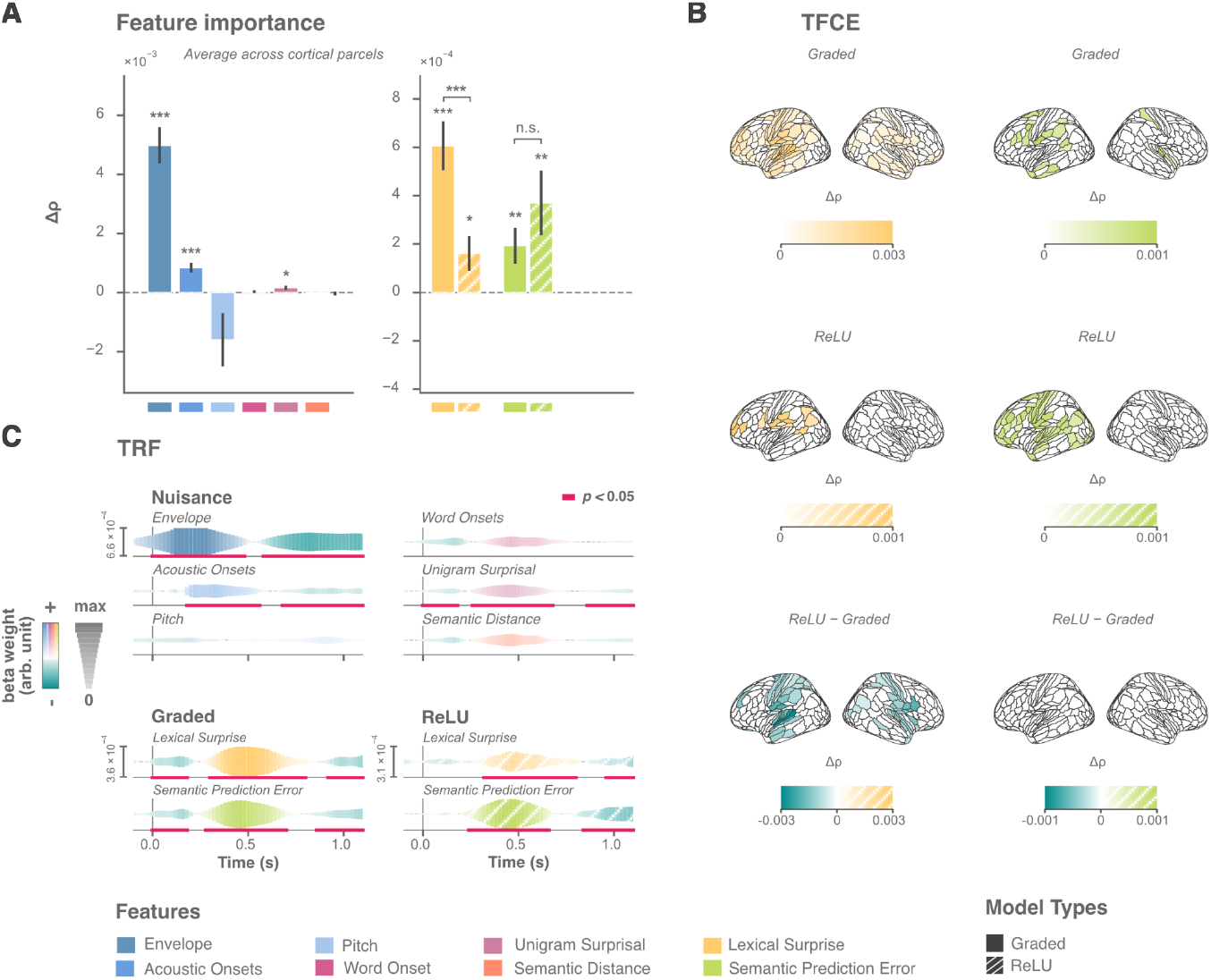
Whole brain cortical tracking. (A)Feature-importance summary across cortical parcels. Each bar shows the mean loss in held-out prediction accuracy (Δρ) after ablating each predictor from the full encoding model, averaged across all 360 cortical parcels. Nuisance predictors (left) enter only in graded form; contextual-prediction features (right) are shown for graded (solid) and rectified-linear-unit (ReLU)-gated (hatched) formulations. Spatial inference for contextual-prediction regressors is shown in (B). (B)Spatial feature-importance maps under threshold-free cluster enhancement (TFCE), with significance assessed against a permutation null at α = .05 (corrected). Both contextual predictors improved cortical encoding under both formulations: lexical surprise was significant across a broadly bilateral peri-Sylvian network, and semantic prediction error across a smaller, predominantly left-lateralised fronto-temporal network. The ReLU - Graded contrast was significantly negative for lexical surprise - the graded model outperformed the gated one over a right-lateralised motor and opercular extent - and showed no significant effect for semantic prediction error (0/360 parcels surviving correction). Peak parcels and cluster extents are descriptive, indicating where effects were most strongly expressed. (C)Group-level TRFs, computed over the parcels showing significant feature importance in (B) and tested against a block-derangement null with cluster-based permutation. Only predictors with a significant feature-importance effect in (B) were submitted to TRF inference; remaining traces are shown for completeness. Nuisance traces reproduced their established morphologies. Contextual predictors are shown under graded (solid) and ReLU (dashed) formulations, with positive deflections over the canonical word-response window and later negativities. Bars and ribbons show group mean ± SEM. Significant TFCE clusters in (B) are shown by coloured parcels; significant TRF windows in (C) are marked by thick pink lines beneath the corresponding trace. Significance levels: \**p* < .05, \*\**p* < .01, \*\*\**p* < .001. *N* = 28.

Group TRFs reproduced canonical morphologies, with positive deflections peaking ∼300-800 ms for lexical surprise and ∼280-700 ms for semantic prediction error, led by earlier (until ∼180 ms) and followed by later (∼950-1200 ms) negativities (Figure 2C). This broad similarity in timecourse is inline with recent findings across hierarchies of processing in both vision ^18^ and audition ^12^, but it is worth noting here that these weights are shaped by regularisation and by correlations between predictors - contextual embeddings in natural language are in themselves correlated so that effects can appear before words are processed even ^19^. The polarity of source-localised responses is also not straightforwardly interpretable - deflection sign reflects a mix of source-estimation and predictor-coding conventions rather than physiology - so while opposite-signed deflections may still index distinct processes, the sign of a deflection does not have a direct physiological interpretation. We therefore test the time courses statistically and read clusters as broad windows indicating where effects are most pronounced, but our inferential claims rest mainly on encoding accuracy (whether a feature improves held-out prediction, Δρ) rather than the detailed shape of the TRF.

This whole-brain result treats the cortex as homogeneous, yet we were interested in also determining differences across the cortical language hierarchy. We thus examined the same feature-importance contrast within four a-priori defined left-hemisphere ROIs (also see Table S1), from early sensory to higher-order cortex: early auditory (Figure 3A-B), temporal-semantic (Figure 3C-D), parietal semantic-integration (Figure 3E-F), and frontal (Figure 3G-H). In early auditory cortex, lexical surprise improved encoding only under the graded model (*W* = 348, *p* < .001, *r*_*rb*_ = 0.71) with a significant advantage over the gated model (*W* = 61, *p* < .001, *r*_*rb*_ = 0.70), whereas semantic prediction error did not reach significance for either formulation and contrast - suggesting a sensory stage tracking the statistical structure of the word stream but not yet meaning-level mismatch. Semantic prediction error improved encoding from the temporal-semantic ROI onward (temporal *t*(27) = 2.63, *p* = .007, *d* = 0.50; parietal *t*(27) = 2.62, *p* = .007, *d* = 0.50; frontal *t*(27) = 1.94, *p* = .031, *d* = 0.37). Across these same regions the graded model’s advantage for lexical surprise diminished: the graded-gated contrast was significant, but numerically weaker, in the temporal (*t*(27) = -3.31, *p* = .003, *d* = 0.62) and parietal (*t*(27) = -2.11, *p* = .044, *d* = 0.40) ROIs, and fell short of significance in frontal cortex (*t*(27) = -1.96, *p* = .060, *d* = 0.37). For semantic prediction error the graded and gated formulations did not differ significantly in any ROI (temporal *W* = 159, *p* = .327; parietal *W* = 182, *p* = .646; frontal *W* = 132, *p* = .109).

**Figure 3.**
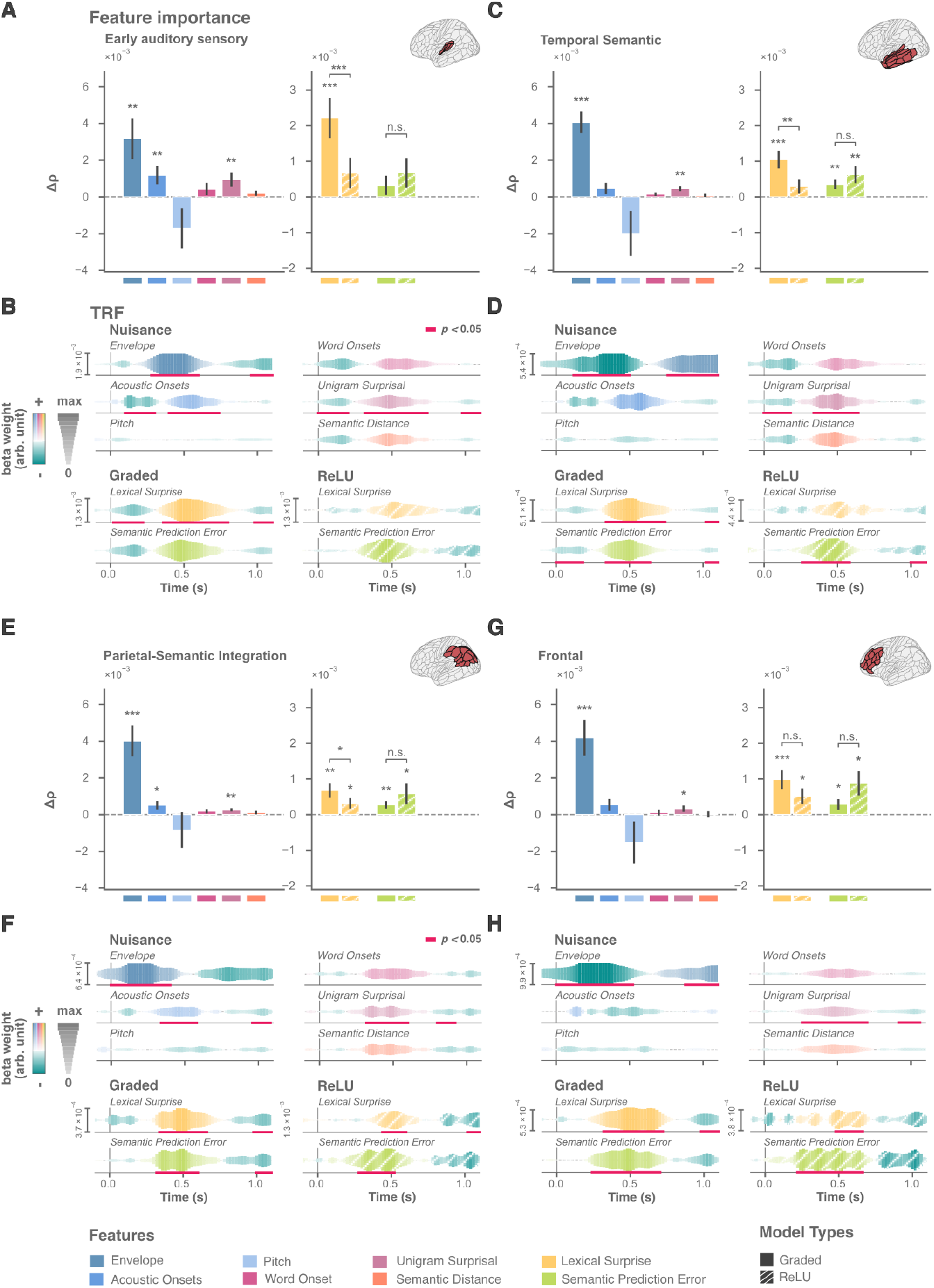
Auditory cortex predominantly tracks lexical surprise in a graded form, with semantic prediction error encoded in later stages and with no graded-gated difference. Feature importance and temporal response functions (TRFs) within four a-priori left-hemisphere ROIs, ordered from early sensory to higher-order cortex: early auditory (A, B), temporal-semantic (C, D), parietal semantic-integration (E, F), and frontal (G, H). ROI extents are shown on the inset cortical surface in each bar panel. (A, C, E, G) Feature importance per ROI. Each bar shows the mean loss in held-out prediction accuracy (Δρ) after ablating a predictor from the full encoding model. Nuisance predictors (left) enter only in graded form; the lexical-surprise and semantic-prediction-error predictors (right) are shown for graded (solid) and ReLU-gated (hatched) formulations. Asterisks above bars denote one-sided tests against zero (Δρ > 0); asterisks above brackets denote two-sided graded−gated contrasts. Tests are one-sample t-tests where subject-level Δρ passed Shapiro–Wilk normality and Wilcoxon signed-rank tests otherwise (see Methods). In no ROI did the ReLU formulation significantly outperform the graded one. Semantic prediction error improved encoding from the temporal-semantic ROI onward but not in early auditory cortex, while the graded model’s advantage for lexical surprise was strongest in early auditory cortex and progressively weaker in higher-order regions. (B, D, F, H) Group-level TRFs per ROI, tested against a block-derangement null with cluster-based permutation. Only predictors with a significant feature-importance effect in the corresponding bar panel were submitted to TRF inference; remaining traces are shown for completeness. Nuisance and contextual predictors are shown under graded (solid) and ReLU (dashed) formulations. Because source-localised responses overlap in time, TRF morphologies are descriptive of how responses distribute across the hierarchy rather than evidence for regionally distinct generators. Bars and ribbons show group mean ± SEM. Significant TRF windows are marked by thick pink lines beneath the corresponding trace. Significance levels: \**p* < .05, \*\**p* < .01, \*\*\**p* < .001. *N* = 28.

The TRF morphologies were consistent with this ascent. Responses to contextual-prediction regressors occupied a common positive window across ROIs (∼250-700 ms); the two features varied with hierarchical level. First, lexical surprise carried an additional early deflection (∼20-220 ms; cluster *p* = .006) most prominent in early auditory cortex and not evident in the higher-order ROIs. Second, later negative components (∼980-1200 ms) emerged for lexical surprise across all four ROIs (cluster *p* ≤ .021), whereas for semantic prediction error a late negativity appeared only in the temporal and parietal ROIs (cluster *p* ≤ .024) and not in early auditory or frontal cortex.

In no ROI did the ReLU formulation outperform the graded one, yet the graded model’s advantage over the gated one was not constant across the hierarchy: strongest in early auditory cortex and progressively weaker in higher-order regions, where for semantic prediction error the two formulations no longer differed. Numerically the pattern was reversed in fact for semantic error. This pattern suggests that cortical encoding may not stay strictly graded in later processing stages, but begins to depart toward a nonlinearity that our thresholding (ReLU) formulation - specified for the hypothesised brainstem gate - captures.

### Brainstem-, but not cerebellar-localised responses, selectively gate semantic prediction error

We next turned to analysing neural responses localised to brainstem structures, as well as the cerebellum as an anatomical control. To sum, the source-localised brainstem ROI showed that gated semantic prediction error outperformed its graded counterpart, while the cerebellar ROI did not.

Specifically, because deep-source reconstruction with MEG warrants caution, the first additional step we took was to examine the brainstem alongside a cerebellar region-of-interest: the cerebellum has been resolved with source-localised MEG during speech ^15^ and is engaged by speech processing more generally ^20^, but it is not part of the neuromodulatory systems indexed by pupil dilation. A brainstem-linked update process should therefore appear in the brainstem but not in this anatomically adjacent control.

For brainstem responses (Figure 4A-B), envelope (*t*(27) = 2.09, *p* = .023, *d* = 0.39) and lexical surprise (*t*(27) = 1.75, *p* = .045, *d* = 0.33) improved encoding under the graded model; no other predictor reached significance (all *p* ≥ .15). Under the gated model, semantic prediction error improved encoding (Wilcoxon *p* = .035, *r*_*rb*_ = 0.39), and the gated-graded contrast was significant for semantic prediction error (Wilcoxon *p* = .036, *r*_*rb*_ = 0.45) but not for lexical surprise (*p* = .17). Thus, while the broad numerical interaction between model and surprise type shares some overlap with the cortical response - that the graded function better describes processing for lexical error but the gated function for semantic error – the brainstem interaction is predominantly driven by a strong response for the thresholded semantic error. Similarities might sometimes reflect some leakage in the localisation functions, but these differences are potentially interesting - especially if they overlap with pupillary responses.

**Figure 4.**
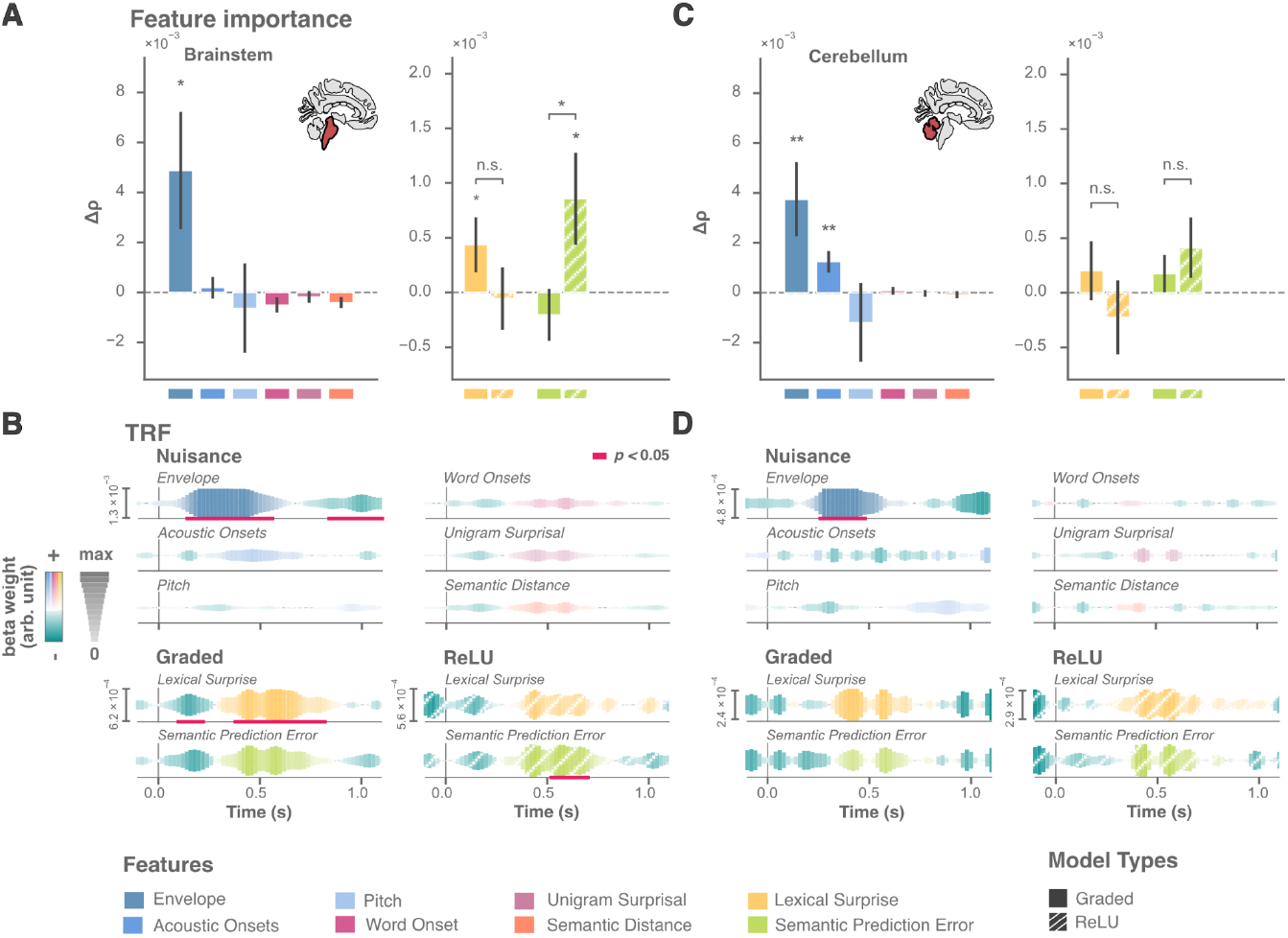
Brainstem, but not cerebellum, selectively gates semantic prediction error. Feature importance and TRFs for the source-localised brainstem (A, B) and cerebellum (C, D) ROIs; ROI extents shown on the inset. (A, C) Feature importance (Δρ) per region. Nuisance predictors (left) enter only in graded form; lexical-surprise and semantic-prediction-error predictors (right) are shown for graded (solid) and ReLU-gated (hatched) formulations. Asterisks above bars denote one-sided tests against zero (Δρ > 0); asterisks above brackets denote two-sided ReLU - Graded contrasts. In the brainstem the contrast was significant for semantic prediction error but not lexical surprise; in the cerebellum neither contextual predictor nor neither contrast was significant. (B, D) Group-level TRFs, tested against a block-derangement null with cluster-based permutation. Only predictors with a significant feature-importance effect in the corresponding bar panel were submitted to TRF inference; remaining traces are shown for completeness. Bars and ribbons show group mean ± SEM. Significant TRF windows are marked by thick pink lines beneath the corresponding trace. Significance levels: \**p* < .05, \*\**p* < .01. *N* = 28.

The TRFs were consistent with this above picture: graded lexical surprise yielded a positive cluster at 380-820 ms (*p* < .001), preceded by an early negativity (100-220 ms; *p* = .032), and gated semantic prediction error produced a positive cluster at 520-700 ms (*p* = .013), alongside the expected early auditory envelope response (140-560 ms, *p* < .001). Notably, the gated semantic brainstem response (∼520-700 ms) fell between the earlier cortical responses to semantic prediction error (∼250 ms onward) and the late cortical components (∼1000-1200 ms), suggesting a brainstem stage interposed between early and late cortical processing.

The cerebellum (Figure 4C-D) showed a different profile, confined to low-level acoustics. Under the graded model, envelope (*t*(27) = 2.53, *p* = .009, *d* = 0.48) and acoustic onsets (*t*(27) = 2.89, *p* = .004, *d* = 0.55) improved encoding; neither contextual predictor reached significance (all *p* ≥ .07), and neither gated-graded contrast was significant (lexical *p* = .13; semantic *p* = .42). The cerebellar TRF showed only an envelope cluster (260-480 ms, *p* =.027).

### Pupil diameter, but not its derivative, mirrors brainstem gating of semantic prediction error

To sum, pupil diameter tracked both forms of surprise, with the same overall pattern as seen in the neural response - numerically greater model fit for a graded function representing lexical surprise and a thresholded function representing semantic surprise. Like with the brainstem-localised signature, it was the improvement for the gated, over graded, semantic function that was particularly robust.

We analysed the pupillary signal in two components: the raw diameter, indexing slower tonic pupil-linked activity, and the first temporal derivative, indexing faster phasic activity more specifically associated with phasic noradrenergic responses ^16,21,22^. To accommodate the approximately 1 s lag between noradrenergic activity and pupil dilation ^16^, the TRF lag window was doubled to 2.4 s. Inference used one-sample tests on Δρ (Student’s t or Wilcoxon signed-rank, selected per predictor; see Methods).

For pupil diameter (Figure 5A-B), no nuisance predictor improved encoding (all *p* ≥ .31). Under the graded model, both lexical surprise (Wilcoxon *p* = .030, *r*_*rb*_ = 0.41) and semantic prediction error (*p* = .025, *r*_*rb*_ = 0.42) contributed; under the gated model, both remained significant, with the larger effect for semantic prediction error (lexical surprise *p* = .039, *r*_*rb*_ = 0.38; semantic prediction error *p* = .006, *r*_*rb*_ = 0.53). Critically, the gated-graded contrast separated the two predictors: for semantic prediction error the gated formulation outperformed the graded one (Wilcoxon *p* = .045, *r*_*rb*_ = 0.43), while for lexical surprise it did not (*p* = .54) - the same selective gating pattern as in the brainstem (noting that the inferential statistics to analyse the commonalities are presented in the next section). The TRFs showed the late clusters expected, given the lag between brainstem and pupillary responses ^16^: graded lexical surprise at 1320-2400 ms (*p* = .022) and semantic prediction error under both formulations (graded 720-2160 ms, *p* = .017; gated 1240-2260 ms, *p* = .030).

**Figure 5.**
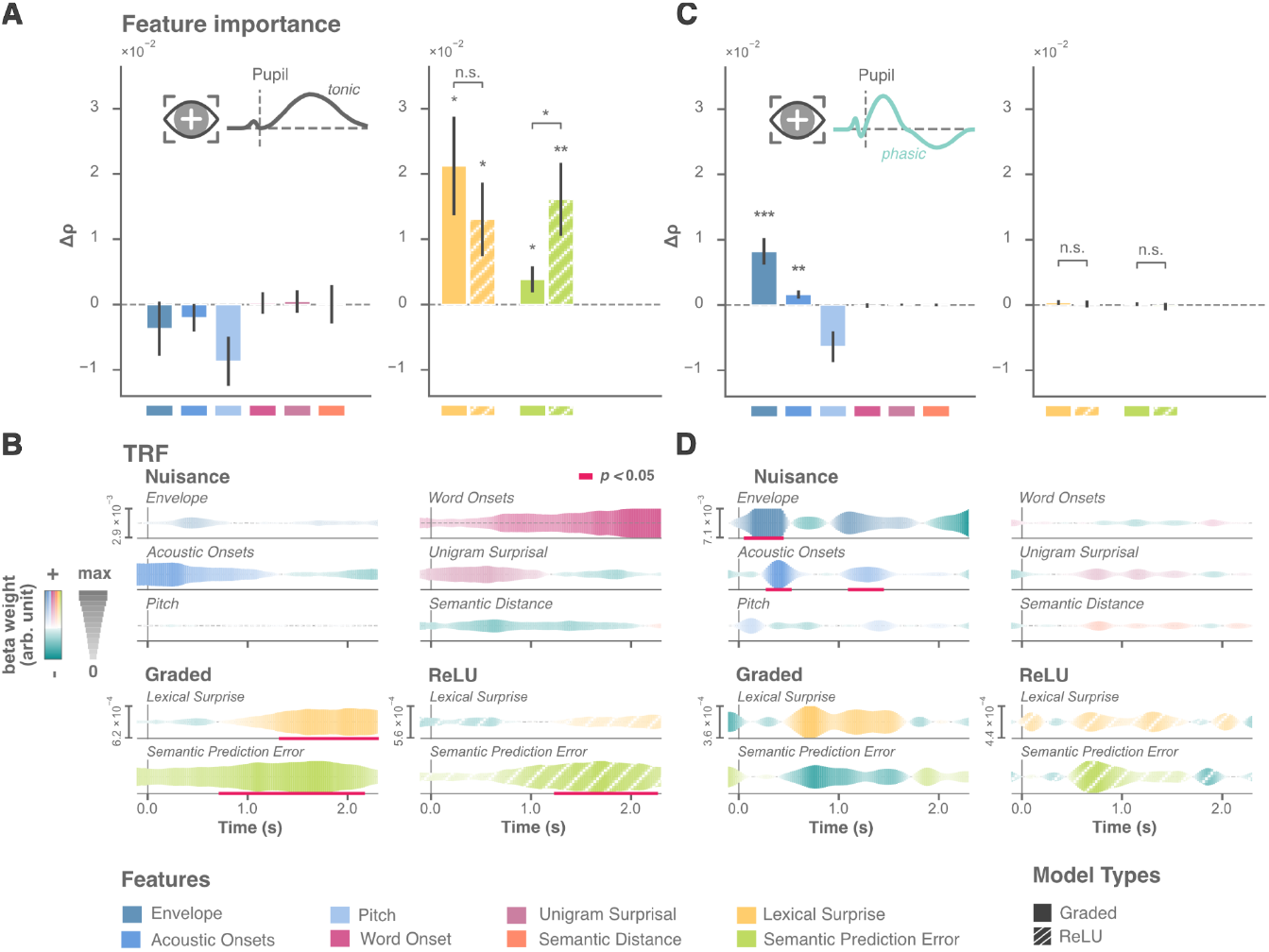
Pupillary response to lexical and semantic prediction error. Feature importance and TRFs for pupil diameter (tonic; A, B) and the pupil derivative (phasic; C, D); pupil-component schematics inset. (A, C) Feature importance (Δρ). Asterisks above bars denote one-sided tests against zero (Δρ > 0); asterisks above brackets denote two-sided ReLU - Graded contrasts. For pupil diameter no nuisance predictor improved encoding, both contextual predictors improved encoding under both formulations, and the contrast was significant for semantic prediction error but not lexical surprise. The derivative was dominated by low-level acoustics, with no contextual effect under either formulation. (B, D) Group-level TRFs, tested against a block-derangement null with cluster-based permutation. Only predictors with a significant feature-importance effect in the corresponding bar panel were submitted to TRF inference; remaining traces shown for completeness. Bars and ribbons show group mean ± SEM. Significant TRF windows are marked by thick pink lines beneath the corresponding trace. Significance levels: \**p* < .05, \*\**p* < .01, \*\*\**p* < .001. N = 28.

The pupil derivative (Figure 5C-D) was dominated by low-level acoustics. Envelope (*t*(27) = 4.06, *p* < .001, *d* = 0.77) and acoustic onsets (*t*(27) = 2.52, *p* = .009, *d* = 0.48) improved encoding; no contextual predictor reached significance under either formulation (all *p* ≥ .12), and neither gated-graded contrast was significant.

### Pupil dilation is specifically coupled to the brainstem-localised, not cortical, response to gated semantic prediction error

As noted, no prior work has examined the pupillary response to contextual predictions during natural speech comprehension, under either the more commonly employed graded formulation or our more novel gated formulation. The fact that the pupil tracked both lexical and semantic surprise, under both formulations, opens a number of questions. Of course our interest in this signature was as a proxy for hypothesised brainstem responses to surprise; the pupil, however, will index not only brainstem activity but a range of cortical and subcortical processes ^16,23^. To determine elements of the pupillary response that may relate specifically to the brainstem response, we implemented a time-resolved mediation analysis ^24,25^. Specifically, this analysis asks where the pupillary signal shares variance with the MEG responses.

Because this mediation rests on the temporal structure of the linguistic TRF in each region, we tested the same four cortical ROIs from the preceding analysis (Figure 3) together with the brainstem ROI. For each region, we quantified pupil-coupling as the reduction in the TRF magnitude when pupil diameter was added as a regressor, evaluated against a block-deranged pupil null in which pupil traces were exchanged across stimulus blocks (Figure 6A-C). This null preserves the temporal structure of the pupillary signal but breaks its alignment with the linguistic input, so a reduction beyond it indicates that the participant’s correctly-aligned pupil dynamics - not a generic pupil-like signal - captured variance shared with the neural response. Inference used 2D cluster-based permutation across TRF lags and pupil shifts, with familywise correction across the predefined ROI × lag × pupil-shift search space (see Methods). We focused on the two predictor formulations that carried the main effects across the preceding analyses: graded lexical surprise and gated semantic prediction error.

**Figure 6.**
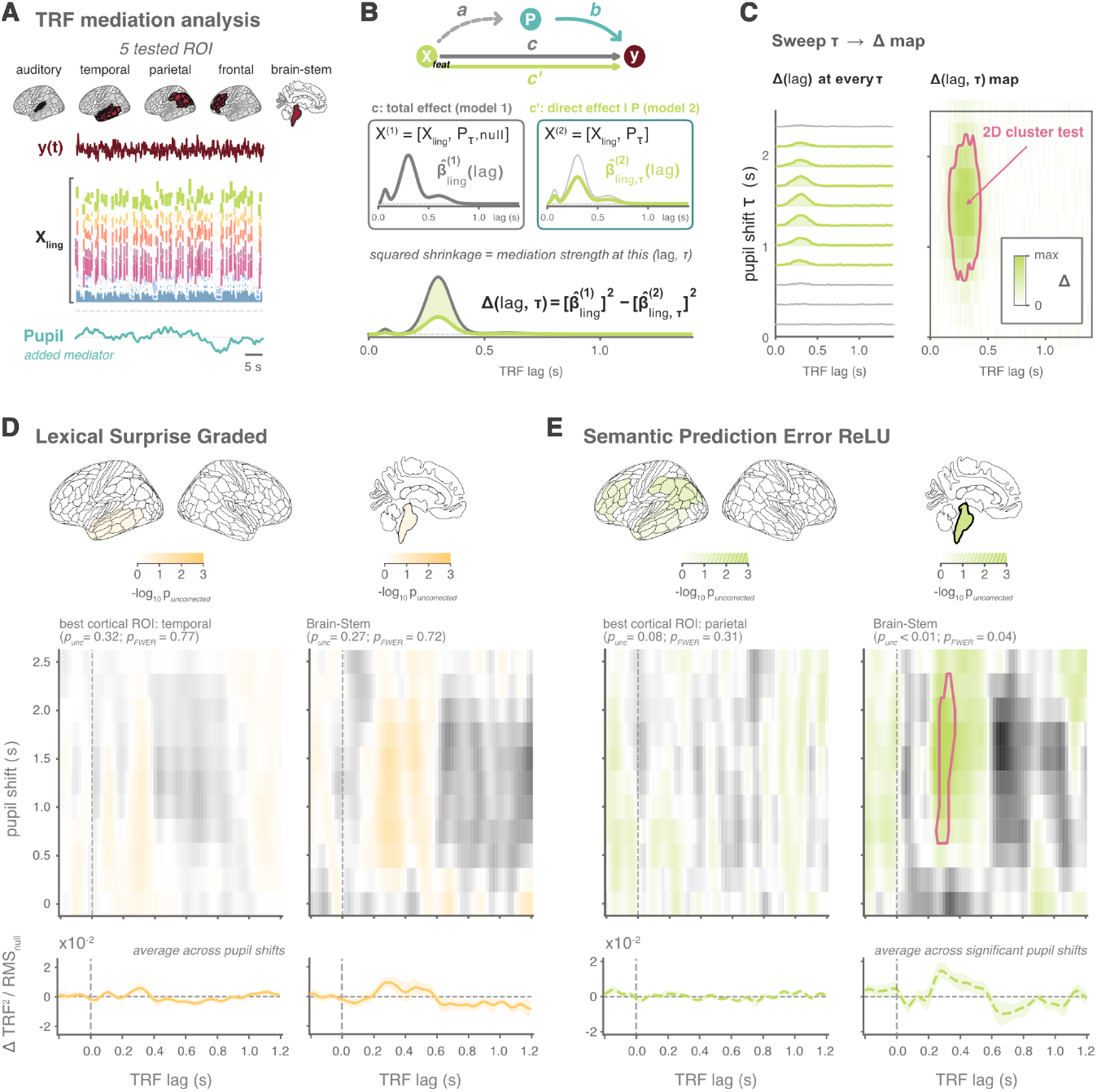
Pupil-linked arousal is specifically coupled to brainstem-localised, not cortical, responses to gated semantic prediction error. A) Analysis setup. Pupil-coupling was tested in five a-priori regions - four left-hemisphere cortical ROIs spanning the language-processing hierarchy (early auditory, temporal-semantic, parietal semantic-integration, frontal) and the brainstem ROI. Traces show a 30-s excerpt from one example subject. (B)Pupil-coupling statistic. For each region, the linguistic TRF was estimated with a correctly-aligned pupil mediator and with a block-deranged pupil null; pupil-coupling is the reduction in squared linguistic TRF between them (see Methods). (C)Pupil shift τ was swept across delays to give a Δ(lag, τ) map over neural TRF lag and pupil shift. Pink outlines mark clusters significant after familywise correction across the ROI × lag × pupil-shift space. (D)Graded lexical surprise. No region showed significant coupling (smallest *p*_*familywise*_ = .72, brainstem; temporal-semantic *p* = .77). Brain maps show per-parcel −log_10_(*p*_*uncorrected*_) for visualization only; inference was at the ROI level. (E)Gated semantic prediction error. The brainstem showed significant coupling (*p*_*familywise*_ = .041; ∼260-360 ms lags, ∼0.75-2.25 s pupil shifts), while no cortical ROI survived correction (strongest, parietal, *p*_*familywise*_ = .31). Colour maps show Δ, the corrected reduction in squared linguistic TRF magnitude. Pink outlines denote significant 2D clusters after familywise correction. *N* = 28.

For graded lexical surprise, no region showed significant pupil-coupling. The strongest cortical effect was in the temporal-semantic ROI (smallest cluster *p*_*uncorrected*_ = .32, *p*_*familywise*_ = .77), and the brainstem ROI similarly did not reach significance (smallest cluster *p*_*uncorrected*_ = .27, *p*_*familywise*_ = .72; Figure 6D). The pupillary effect of graded lexical surprise seen in the encoding analysis is therefore not traceable to any of the tested regions.

For gated semantic prediction error, the pattern differed sharply. Cortical ROIs again failed to reach significance after correction, the strongest being the parietal semantic-integration ROI (cluster *p*_*uncorrected*_ = .08, *p*_*familywise*_ = .31). The brainstem ROI, by contrast, showed significant pupil-coupling (cluster *p*_*uncorrected*_ = .006, *p*_*familywise*_ = .041), over a cluster spanning neural TRF lags of approximately 260-360 ms and pupil shifts of approximately 0.75-2.25 s (peak *t* = 3.50, *d* = 0.66; Figure 6E). The lag-by-shift geometry indicates that the brainstem response around 260-360 ms was coupled to the pupillary response approximately 0.75-2.25 s later - consistent with the established ∼1 s lag between phasic noradrenergic activity and pupil dilation ^16^, propagated through the slower tonic component captured by pupil diameter.

The mediation thus identifies a doubly selective coupling: the pupil-brainstem link is specific to gated semantic prediction error (not graded lexical surprise), and to the brainstem ROI (not to any tested cortical region, including those that showed encoding effects). This specificity strengthens the idea that the (noisy) brainstem-localised and (indirect) pupil-diameter signatures of gated semantic prediction error reflect a common process.

## Discussion

We here demonstrate how the brain and pupil track contextual predictions during natural speech comprehension. Across analyses, lexical surprise was best described by a graded response, whereas semantic prediction error was better captured by a thresholded (gated) one, where the brain and pupil respond only once the error exceeds a recent contextual baseline. There were also interesting differences in the relative strength of these effects across measures. The auditory cortex tracked LLM-derived lexical surprise in a graded, continuous manner, with sensitivity to semantic error emerging only in later temporal, parietal, and frontal responses. By contrast, source-localised brainstem responses showed the gated signature most strongly, with the thresholded model outperforming the graded one specifically for semantic surprise. The pupil mirrored this response to gated semantic error and a pupil-coupling analysis showed that variance in the pupillary response was shared with the brainstem-localised, but not cortically-localised, signature. Together, these results suggest that the auditory cortex maintains a high-fidelity, continuous map of predictive lexical information, whereas a pupil-linked brainstem system acts as a selective gate, engaged specifically by meaning-disrupting events that signal the need to update internal models of the world.

The cortical encoding observed here builds on a now-substantial literature fractionating prediction representation across the cortical hierarchy ^11–13^. We show that, broadly speaking, the cortical response to error is better modelled as a graded function, by comparing this typically-employed function against a gated one. This graded picture turned out to be not, however, entirely uniform. Resolving the cortical hierarchy into regions of interest, the graded model’s advantage for lexical surprise was largest in early auditory cortex and weakened toward higher-order regions; for semantic prediction error the gated formulation was numerically favoured across the cortex, though the contrast reached significance in no region. This pattern of the graded model losing superiority toward higher order areas is arguably a function of downstream influences of the gating functions reflected in brainstem-localised and pupillary processing. Notably however, the cortical response did not show much sign of mediation by the pupillary signature, so the neural responses are likely reflecting complex signatures by these higher order structures. For example, some recent work suggests that we may compute subjective or cognitive prediction errors, which may be only partly related to their objective counterparts ^26,27^.

A thresholded response to semantic prediction error - observed across the brainstem-localised neural response and the pupillary signature - suggests that meaning-related mismatches may invoke responses thought to play a role in model-updating. The specifics of this finding require some careful consideration. Notably, the effect was carried by raw pupil diameter rather than its derivative, which may at first seem at odds with canonical accounts emphasising phasic locus coeruleus bursts ^17,28^. However, meaning-relevant prediction errors unfold against a context that builds across seconds, and given the ∼1 s lag between phasic noradrenergic activity and pupil dilation ^16^, the late TRF windows we observe (peaking ∼720-2160 ms) are consistent with summed phasic activity expressed in the slower diameter signal rather than a categorically tonic process. These findings build on work showing that pupil measures track stimulus-derived acoustic and lexical features during naturalistic narrative listening ^29–31^, and that pupil responses can index Bayesian belief updating, decision uncertainty, and surprise ^10,21,22,32–34^, including surprisal in dynamic auditory sequences ^35–37^. Urai and colleagues ^21^ further showed that pupil-linked arousal tracks decision uncertainty in the moment and predicts how that uncertainty biases subsequent choices - implicating arousal in signals that propagate forward rather than merely registering the current state, paralleling the selective reflection of threshold-crossing semantic mismatch we identify here for naturalistic language. We extend this work by showing a pupillary response to surprise in natural language comprehension.

The convergent signatures across tonic pupil diameter, source-localised brainstem responses, and the pupil-coupling analysis is a central contribution of this work. Each measure alone would invite much caution: MEG source estimates of brainstem activity are methodologically demanding, and pupil responses are slow and noisy and reflective of a range of processes. However, observing this convergence across measures is particularly encouraging, and opens new avenues for future research. The inference’s force comes from independent measures converging on the same conclusion - the gated formulation outperformed the graded one in pupil diameter, in the brainstem, and in the pupil-coupling analysis, selectively for semantic prediction error. The cerebellum, an anatomically nearby deep source implicated in temporal prediction during speech ^15^ and recently shown to contain language-selective regions ^20^, showed no such signature: its source estimates were dominated by acoustic structure, with no contribution from either contextual predictor under either formulation. The brainstem gating effect therefore does not appear to be a generic subcortical artefact but anatomically and computationally selective, fitting accounts in which neuromodulatory systems implement threshold-dependent gating of model updating rather than continuously tracking prediction error ^14^.

Two features of this coupling invite tentative comment. First, the pupil-coupled brainstem response was expressed at an earlier latency (∼260-360 ms) than the gated semantic cluster that dominated the brainstem encoding estimate (∼520-700 ms), though the descriptive nature of cluster bounds precludes treating these as cleanly separable windows. Because the mediation analysis indexes shared variance rather than raw amplitude, it may foreground a different, lower-amplitude part of the brainstem response than the encoding estimate, such that the two windows need not reflect distinct generators. That this brainstem response shares variance with pupillary effects is consistent with a noradrenergic contribution, given that pupil dilation is driven substantially by the locus coeruleus ^16,17^. Second, of course our brainstem ROI does not resolve individual nuclei. With an MEG signal this separation would seem questionable at present, but a range of nuclei, in addition to locus coeruleus, could in principle drive responses. Importantly however, the mediation analysis does show that there is a component of this signal which couples to the pupillary one, and therefore this coupled variance would arguably need to be generated by a structure whose activation is also reflected in pupil dilation.

The difference between semantic and lexical surprise in the shape of function is striking and, we suggest, principled. Not all prediction errors are equivalent in relevance. Lexical surprise primarily indexes the moment-to-moment statistical structure of the unfolding word stream, and tracking it continuously is arguably necessary for temporal parsing, segmentation, and the maintenance of an ongoing scaffold over linguistic input - gating it would dissolve that scaffold. Semantic prediction error, by contrast, indexes mismatch in meaning, and the kind of signal that should warrant updating one’s situational or conceptual model, but only perhaps when the mismatch is large enough to be worth the cost of revision. Which error signals become gated is therefore, we assume, not arbitrary: signals whose function is to maintain a continuously updated parse should be tracked continuously, while signals whose function is to flag the need for model revision should be selectively rectified.

These findings have implications for predictive processing theory beyond language. Most accounts treat prediction error as a single quantity driving both perceptual updating and learning; our data suggest these may be computationally distinct operations in dissociable systems. The sensory cortex may maintain a continuous representation of prediction error across its dynamic range - the signal needed to support graded inference and online interpretation - while the pupil-linked brainstem system operates more like a thresholded gate, engaged when error magnitude exceeds a context-sensitive baseline and only when that error reflects behaviourally-relevant mismatch. A recent synthesis ^38^ noted that subcortical and neuromodulatory contributions to prediction-error signalling have been characterised primarily in reward and decision-making, while sensory prediction errors have been studied almost exclusively in cortex. Recent work in mice begins to bridge this gap, showing that dorsal striatal dopamine encodes sensory alongside reward prediction errors and causally biases subsequent perceptual choices ^39^. Our findings extend this convergence to a different neuromodulatory axis and to continuous naturalistic perception in humans, where prediction error arises without explicit trial-level reward and the gated signature is selective for meaning-disrupting events - pointing to a brainstem-linked neuromodulatory architecture that may generalise across species, neuromodulators, and modalities. The question is then not whether prediction error is graded or thresholded, but at which level of the system each computation occurs. The relative timing of our signatures, while tentatively interpreted, is consistent with such a layered architecture: the gated brainstem response started later than the auditory cortical response to graded lexical error, but preceded some gated cortical responses in higher-order structures. Continuous error representations may thus support moment-to-moment inferential dynamics, while selective neuromodulatory gates determine when those representations cross the threshold of engaging model-updating - an architecture echoing proposals from perceptual learning ^14^ and recent within-cortex dissections of uncertainty ^40^.

MEG source localisation of brainstem activity should be treated with caution, and the strength of the present inference lies in its convergence with the independent pupillary signal. The pupil-coupling analysis identifies shared variance rather than causal direction of course, leaving open the mechanism by which pupil-linked responses and brainstem activity become coordinated. The ReLU is one specific formalisation of thresholded gating, chosen for its simplicity and centrality to neural-network architectures; it provides a useful computational vocabulary bridging predictive-processing accounts and machine-learning models of language, but future work could establish whether it best captures the relevant nonlinearities. Participants were also informed that they would be questioned on story content in our study, which may have directed attention toward semantic processing and amplified the semantic-specific effect; the dissociation should be tested also under fully passive listening. Because we analysed LLM-derived surprise from naturalistic input, cleanly dissociating predictive from other contextual signals is difficult ^19^, and lab-based manipulations of probability could seek to replicate these patterns. Finally, the present results derive from a single language, modality, and LLM; cross-linguistic and cross-modal replication will test the architecture’s generality. Probing brainstem neuromodulation more directly remains challenging in naturalistic paradigms: 7T fMRI and neuromelanin-sensitive imaging of these structures offer spatial specificity at the cost of temporal resolution and acoustic compatibility - and could adjudicate, for example, whether a locus coeruleus component underlies the coupling described above - while pharmacological manipulation could provide further convergence ^41^.

In conclusion, we here show that the brain and pupil track lexical and semantic surprise when listening to a story, albeit not in precisely the same fashion. Broadly across all analyses, lexical deviations elicited a more graded response, whereas semantic surprise only elicited responses when crossing a threshold level of error. There were also some interesting distinctions in the relative strength of these effects across the brain and pupil. The auditory cortex continuously tracks lexical prediction error in a graded manner, while a pupil-linked brainstem system selectively engages when meaning-relevant errors exceed a recent baseline. This division of labour - continuous representation of lexical deviations in the auditory cortex alongside selective gating according to meaning in the brainstem-pupil axis - suggests that the perception-learning interface during natural language comprehension is implemented not as a single error signal but as a coordinated architecture in which cortical graded errors provide information to a subcortical gating mechanism, with subsequent downstream influences across cortical hierarchies.

## Methods

### Participants

Twenty-eight adults participated in the experiment (final sample after exclusion criteria; see below). All participants were native German speakers, provided written informed consent, and reported normal hearing, normal or corrected-to-normal vision, and no history of neurological or psychiatric conditions. They were compensated with €10 per hour or course credit. The experiment was conducted in accordance with the Declaration of Helsinki and approved by the Ethics Committee of the University of Salzburg. Participants were excluded from analysis if more than 50% of pupil samples were lost to blinks across the listening blocks. This was the case for one participant in the sample.

### Stimuli and experimental design

Before the experiment, participants’ head shapes were digitised using a Polhemus Fastrak (Polhemus, Colchester, VT) by recording cardinal landmarks (nasion and pre-auricular points) and approximately 300 additional points distributed across the scalp.

Participants listened to four German-language short stories, each narrated by either a male or a female speaker and presented in a separate block of approximately 3-4 min (mean ± SD: 2,065 ± 32 words per participant; range: 2,000-2,123). Stimuli were delivered binaurally at 40 dB above the individual hearing threshold via pneumatic headphones. Participants were instructed to attend to the speaker and direct their gaze towards a central black cross on the screen. After each block, they answered true/false comprehension questions (e.g., “Das Haus, in dem Sofie lebt, ist rot” - “The house Sofie lives in is red”) and rated their task engagement and perceived difficulty on a 5-point Likert scale. The experiment was implemented in Psychtoolbox-3 ^42,43^ using the Objective Psychophysics Toolbox ^44^.

### Data acquisition

#### MEG

MEG was recorded using a whole-head Elekta Neuromag Triux system (Elekta Oy, Helsinki, Finland) housed in a passively shielded room (AK3b, Vacuumschmelze, Hanau, Germany), with 102 magnetometers and 204 planar gradiometers (102 sensor positions) at a 1 kHz sampling rate (hardware filters 0.1-330 Hz). External noise was suppressed and inter-block head positions were realigned using the signal-space-separation algorithm implemented in MaxFilter (version 2.2.15; Elekta Oy).

#### Eye-tracking

Binocular eye-tracking was performed with a TRACKPixx3 system (VPixx Technologies, Saint-Bruno, Canada) equipped with a 50 mm lens, sampling at 2 kHz. Recordings began after a 13-point calibration and validation procedure. Participants were seated 82 cm from the screen with their head stabilised by a chin-rest. Stimulus-onset triggers from the experimental control software were sent in parallel to the MEG and TRACKPixx systems, yielding sample-accurate temporal alignment between the two recordings. Gaze position (horizontal, vertical) and pupil diameter were saved natively by the TRACKPixx system.

### Preprocessing

#### MEG

MEG preprocessing was carried out in MNE-Python ^45^ using the AlmKanal M/EEG preprocessing pipeline developed at the Salzburg Brain Dynamics Lab (https://github.com/schmidtfa/AlmKanal). For each participant, the four blocks were loaded, parsed into trials based on stimulus-onset triggers, and segments outside experimental trials were marked as bad. Continuous concatenated data were submitted to MaxFilter-based MultiBlockMaxwell processing to harmonise head positions across blocks, followed by independent component analysis (ICA) using the Picard algorithm ^46^ on data low-pass filtered at 100 Hz and resampled to 200 Hz. Cardiac and ocular components were identified by correlation with the ECG and EOG channels, respectively (correlation threshold = 0.3, training threshold = 3 z-scores), and removed. This automated procedure identified a median of 2 artefactual components per participant for removal, range = 1-4 (*n* = 4 and *n* = 2 at the minimum and maximum, respectively). The continuous data were then band-pass filtered between 0.5 and 8 Hz (zero-phase FIR), the ICA solution was applied, and the cleaned data were time-corrected for the 16.5 ms acoustic delay introduced by the pneumatic headphones and resampled to 50 Hz for subsequent source reconstruction.

#### Eye-tracking

Eye-tracking data were preprocessed in MATLAB (R2025a) using FieldTrip ^47^. Throughout the analysis, data from the right eye were used. Blinks were detected from the pupil signal using the noise-based detection algorithm of ^48^. To remove residual artefacts at blink edges, the surrounding 50 ms for gaze channels and 150 ms for the pupil channel were additionally marked as missing. Gaze coordinates were converted from pixels to degrees of visual angle using the screen geometry (61 × 34.3 cm at 82 cm viewing distance). Saccades were detected using a velocity-based algorithm ^49–52^ implemented on the two-dimensional gaze-velocity profile with a 7-sample smoothing window. Saccade onsets and offsets were identified when velocity exceeded and subsequently fell below five times the median velocity, with a minimum 100 ms inter-saccade interval. Saccades were classified as microsaccades when their amplitude was between 0.057° and 1°, and as macrosaccades when their amplitude exceeded 1°. Gaze coordinates were median-centred. Pupil traces were gap-interpolated linearly, low-pass filtered at 6 Hz using a zero-phase FIR filter, and detrended. Pupil and gaze traces were time-corrected for the 16.5 ms acoustic delay introduced by the pneumatic headphones, downsampled to 50 Hz to match the MEG sampling rate after source reconstruction, and trimmed to the duration of each speech segment. Trials in which more than 50% of pupil samples were missing led to exclusion of the participant (see Participants).

As a general quality and compliance check, we quantified viewing behaviour during listening. Participants made on average 0.26 ± 0.17 blinks/s, 0.96 ± 0.88 microsaccades/s, and 0.32 ± 0.41 macrosaccades/s. The low macrosaccade rate is consistent with predominantly stable fixation during auditory-only stimulation.

#### Pupil residualisation

Blinks and saccades elicit large, seconds-long pupil responses that contribute substantial variance to the pupil signal and can be removed by simultaneous nuisance regression ^53^; gaze position likewise influences measured pupil size and can be accounted for by including it as a regressor rather than by pre-processing correction ^54^. To isolate variance in pupil diameter not attributable to these peripheral oculomotor contributions, we regressed gaze position, blink onsets, and saccade onsets out of the pupil signal using temporal response functions (TRF; ^55–57^). For each block, we fit a ridge-regularised TRF with a window from − 2 to + 6 s relative to each predictor sample. The ridge parameter α was selected within block by three-fold cross-validation over ten log-spaced values, α ∈ {10^1^, •, 10^10^ }, using z-scored predictors and a z-scored pupil response. The estimated oculomotor TRFs (Figure S1) recovered biphasic blink- and saccade-related profiles consistent with those previously reported ^53^. The predicted artefact time course was then subtracted from the observed pupil signal in original units. The residualised pupil diameter was used for all subsequent analyses. The first temporal derivative of the residualised pupil (“pupil derivative”) was additionally computed by first differencing and was subsequently low-pass filtered at 2 Hz ^22^; this complementary signal indexes phasic pupil-linked activity ^16^.

### Source modelling

To recover MEG activity from cortical and subcortical sources, we constructed a mixed source space combining a high-resolution cortical surface with subcortical volumetric sources. Because individual structural MRIs were not available, we used the FreeSurfer ^58^ *fsaverage* template MRI as a common anatomical reference. Co-registration to each participant’s digitised head shape was performed using an automated MRI-to-headshape alignment procedure based on the digitised cardinal landmarks and scalp points.

The mixed source space comprised (i) a cortical surface source space defined on the white-matter surface with oct6 spacing (4,098 sources per hemisphere), and (ii) a volumetric source space sampled at 4 mm spacing within four subcortical/cerebellar labels from the FreeSurfer *aseg* parcellation: Brainstem, Left-Cerebellum-Cortex, Right-Cerebellum-Cortex. A three-shell boundary-element model (5120–5120–5120; inner skull / outer skull / scalp; conductivity values fixed to the MNE defaults of 0.3, 0.006, and 0.3 S/m) was constructed from the template MRI, and the forward solution was computed using mne.make_forward_solution with a minimum source-sensor distance of 5 mm.

A linearly constrained minimum-variance (LCMV) beamformer ^59^ was applied to reconstruct source time courses from the continuous preprocessed MEG. Data covariance was estimated from the full preprocessed signal; noise covariance was estimated from the temporally nearest empty-room recording for each participant. Beamformer weights were applied to the continuous data using mne.beamformer.apply_lcmv_raw, yielding source time courses at every cortical-surface vertex and every subcortical-volume voxel.

Cortical source activity was parcellated into the 360 regions of the *HCP-MMP1*.*0 atlas* ^60^ using mean-flip extraction (mne.extract_label_time_course, mode = mean_flip), which sign-aligns vertex orientations within each parcel before averaging to mitigate orientation-induced cancellation. For each subcortical/cerebellar label, source time courses across the contained voxels were averaged using the mean, because mean-flip extraction is not applicable to volume sources. The Left- and Right-Cerebellum-Cortex parcels were combined into a single bilateral cerebellum signal for analysis. After source reconstruction, parcel time series were segmented into the per-block listening segments and aligned with the corresponding pupil and predictor time series.

### Predictors

All linguistic and acoustic features were aligned to a common 50 Hz time grid matching the neural and pupil signals. Word timing information was obtained from forced alignment of the audiobook transcripts to the speech recordings using the Munich Automatic Segmentation System (MAUS; ^61^). All word-level features were placed as impulses at the corresponding word-onset sample of the speech segment.

#### Acoustic features

Two low-level acoustic predictors were derived from gammatone spectrograms of each speech segment (128 frequency bands, 80-15,000 Hz, equivalent-rectangular-bandwidth spaced, sampled at 1 kHz; ^62^) computed in Eelbrain ^63^. The one-band envelope was obtained by summing the log-transformed (natural log of value plus one) gammatone spectrogram across frequencies. Acoustic onsets were derived by applying a neurally inspired auditory-edge-detection transform (^64^; TRF-Tools, github.com/christianbrodbeck/TRF-Tools; saturation factor *c* = 30) to the log-spectrogram and summing across frequency bands. Pitch was extracted from the raw waveform using Praat’s autocorrelation algorithm via Parselmouth ^65^ at 1 ms resolution, with *F*_0_ values constrained to 0-500 Hz and outliers (|*z*| > 3) replaced by linear interpolation. Each acoustic feature was scaled to [0, 1] within block to remove between-block amplitude differences before alignment to the neural sampling grid.

#### Non-contextual lexical features

Three predictors captured word-level processing components independent of the broader linguistic context. Word onsets were modelled as a binary impulse at each word’s onset sample, controlling for stimulus-locked responses to word presentation per se. Unigram surprisal quantified the rarity of each word according to the German word-frequency distribution and was computed for word *w*_*i*_ as

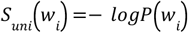

where *P*(*w*_*i*_) was obtained from the wordfreq Python library ^66^. Semantic dissimilarity quantified the semantic distance of each word from its immediately preceding context ^67^. For each word *w*_*i*_ with *fastText* embedding 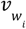 (^68^; German cc.de.300 vectors), semantic dissimilarity was computed as

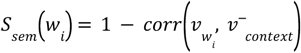

where 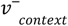 is the average fastText vector across all words preceding *w*_*i*_ within its sentence; for sentence-initial words, the context was taken as the mean vector of the preceding sentence.

#### Contextual prediction features

Two contextual predictors quantified the degree to which each word violated context-conditioned predictions, derived from a pretrained German GPT-2 model (dbmdz/german-gpt2; ^69^) loaded via Hugging Face Transformers ^70^. For each word, the model was conditioned on the unbroken preceding context within the block. Because GPT-2 operates on sub-word tokens, word-level probabilities were obtained by multiplying the conditional probabilities of constituent tokens. Lexical surprise was defined as the negative log-probability of the observed word *w*_*i*_ given its context *w*_1:*i*−1_:

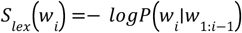

Semantic prediction error quantified the cosine distance between the embedding of the word that actually occurred and the model’s expected semantic vector, following ^12^. For each word boundary, we identified the set of top-*k* most probable next-word candidates, with *k* defined dynamically as the smallest set whose cumulative probability reached 0.90, subject to a minimum of *k* = 40. Candidate tokens were restricted to word-start tokens, excluding pure-whitespace and punctuation-only tokens. For each candidate *c* with fastText embedding *v*_*c*_ and re-normalised probability 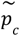, the expected semantic vector was computed as a probability-weighted average:

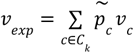

Semantic prediction error for the observed word *w*_*i*_, with embedding *v*_*obs*_, was then defined as the cosine distance to the expected vector:

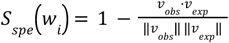

#### ReLU (gated) transformation

To test whether neural and pupillary responses are better captured by a thresholded gate than by a graded representation, we constructed rectified versions of each contextual predictor. *x*_*i*_ denotes the word-level value of a contextual predictor, either *S*_*lex*_ or *S*_*spe*_, at word *i* within a block, *i* = 1, •, *N*. For each block independently, we computed a causal, history-dependent z-score relative to all preceding words in the same block,

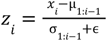

with ϵ = 10^−9^ for numerical stability and *z*_1_ ≡ 0 for the first word in each block, which has no history. The ReLU-transformed predictor was then defined as

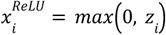

following the rectified-linear-unit nonlinearity widely used in modern neural-network architectures ^71^. Only positive deviations above the local contextual baseline pass through; sub-baseline values are rectified to zero. This transform provides a simple, falsifiable instantiation of thresholded model-updating ^14^. The graded *x*_*i*_ and gated 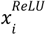 versions of each contextual predictor were entered into otherwise identical encoding models.

### Encoding models

For each participant and each response channel (cortical parcel, brainstem parcel, bilateral cerebellum parcel, residualised pupil diameter, or pupil derivative), we fit TRF encoding models ^55–57^ predicting the response from a set of time-lagged predictors. Letting *y*(*t*) denote the response at time *t* and *S*_*f*_ (*t*) the *f*-th predictor, the encoding model takes the form

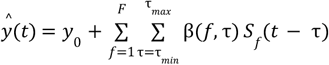

where β(*f*, τ) are the TRF weights for predictor *f* at lag τ, and *y*_0_ is an intercept. The lag window was [τ_*min*_, τ_*max*_] = [− 0. 2, 1. 2] s for MEG responses and [− 0. 2, 2. 4] s for both pupil diameter and the pupil derivative. Within each cross-validation fold, predictors *S*_*f*_ were scaled using StandardScaler(with_mean=False), fit on the training blocks and applied to the held-out block, thereby preserving the sparse impulse structure of word-level features. Responses *y* were z-scored within fold using StandardScaler, again fit on the training blocks and applied to the held-out block.

Two predictor sets were defined. The graded set,

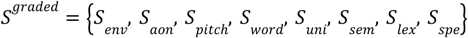

included both contextual predictors in their continuous form. The gated set *S*^gated^ was identical to the graded set except that the two contextual predictors *S*_*lex*_ and *S*_*spe*_ were replaced by their ReLU-transformed counterparts 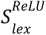 and 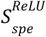; all nuisance predictors (acoustic features, word onsets, unigram surprisal, and semantic dissimilarity) remained graded in both.

TRF weights β were estimated by ridge regression,

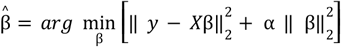

where *X* is the time-lag-embedded predictor matrix. Model performance was evaluated using leave-one-block-out cross-validation across the four speech blocks, implemented with GroupKFold(n_splits = 4). In each fold, models were trained on three blocks and tested on the held-out block. Within each training fold, the ridge parameter α was selected separately for each response channel using RidgeCV with per-target alpha selection from 20 log-spaced values, α ∈ {10^1^, •, 10^10^ }. Predicted and observed time courses were Spearman-correlated across samples within the valid TRF window, yielding a per-channel held-out encoding score ρ. Per-subject ρ values were Fisher-z transformed,

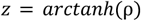

and averaged across folds before group-level inference.

#### Feature importance

To quantify the unique contribution of each predictor to encoding performance, we used a single-feature ablation procedure (as in e.g. ^72–74^). For the graded model, ablations were computed for all predictors. For the gated model, ablations were restricted to the two contextual predictors, because nuisance predictors were identical across graded and gated formulations. Feature importance was defined as the loss in held-out Fisher-z-transformed correlation,

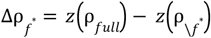

with ρ_*full*_ and 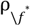 denoting the held-out Spearman correlations obtained from the full and reduced models, respectively. Larger Δρ values indicate that the ablated feature carried unique variance not captured by the remaining predictors. For the contextual predictors, ablation was performed within each model type (graded or gated) so that Δρ values are directly comparable across formulations. We additionally computed the Gated − Graded contrast in feature importance,

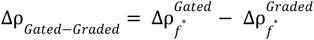

which tests whether the gated formulation captures additional variance over and above the graded one.

#### TRF weights and block-derangement null

For visualisation and statistical testing of TRF time courses, we extracted per-subject TRF weights at a single, globally fixed ridge parameter 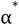, defined as the modal best-α value across subjects, folds, and channels obtained from the full (non-ablated) encoding models. Using a single 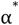 ensures that TRF weights are quantitatively comparable across subjects and models. For each fold, alongside the real-data TRF 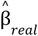, we additionally computed a block-derangement null TRF. For every derangement π of block indices, defined as a permutation with no fixed points, predictor blocks were paired with mismatched response blocks, *S*^π(*b*)^ paired with *y*^(*b*)^, the model was re-fit, and the resulting null TRFs were averaged across derangements,

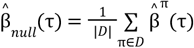

where *D* is the set of derangements of block indices. For four blocks, |*D*| = 9. This null preserves the within-block temporal structure of the predictors while breaking their alignment with the observed neural or pupillary response, providing a matched chance-level baseline against which the real TRF time course was tested.

### Mediation (Pupil-coupling) analysis

To test whether stimulus-locked neural responses to contextual surprise were statistically shared with pupil-linked responses, we asked whether including pupil diameter as an additional regressor reduced the magnitude of the linguistic TRF, relative to a block-deranged pupil null. We refer to this corrected reduction as pupil-coupling. The analysis was restricted to a fixed, a-priori set of regions of interest (ROIs) spanning the language-processing hierarchy (for the complete parcel composition of each ROI, see Table S1); ROIs were defined independently of the present TFCE feature-importance maps. Five ROIs were tested: four left-hemisphere cortical ROIs (early auditory, temporal semantic, parietal semantic integration, and frontal) and the brainstem volumetric parcel. Cortical-ROI surfaces were averaged across constituent parcels within subject before inference.

#### Models

For each subject, ROI, and cross-validation fold, we fit two encoding models. The baseline linguistic-only model recovered the linguistic TRF β_*ling*_ as in the main encoding analysis,

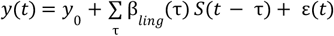

The mediated model additionally included residualised pupil diameter *P*(*t*) at a temporal shift τ_*P*_ ∈ [0, 2. 5] s, sampled in 0.25 s steps,

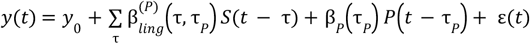

The linguistic TRF in the mediated model, 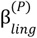, is expected to be smaller in magnitude to the extent that the linguistic response and the lag-shifted pupil response share variance. To prevent edge effects when shifting pupil traces, MEG and predictor time series were cropped by the maximum shift; the pupil was indexed from the corresponding shifted position within the original uncropped block.

#### Pupil-coupling statistic

For each ROI, fold, and pupil shift τ_*P*_, we quantified pupil-coupling at every TRF lag τ as the difference in squared linguistic TRF magnitude between models in which the linguistic regressors were paired with the block-deranged pupil and with the correctly aligned pupil,

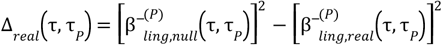

Here, 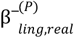 denotes the fold-averaged linguistic TRF after including the correctly aligned pupil trace, whereas 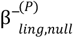 denotes the corresponding TRF after including the block-deranged pupil trace. Positive values therefore indicate that the correctly aligned pupil trace reduced the linguistic TRF more than a temporally structured but block-mismatched pupil trace. We used squared rather than absolute coefficients because (i) squaring is variance-stabilising for Gaussian regression coefficients, (ii) β^2^ is the per-coefficient contribution to explained variance, and (iii) the sign-agnostic property is appropriate when aggregating across parcels whose TRF polarity may differ. Δ_*real*_ (τ, τ_*p*_) was averaged across constituent parcels within each cortical ROI to yield a single 2D surface per subject and ROI over the lag × pupil-shift grid.

#### Block-deranged pupil null

The null component of, 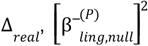, was constructed by re-fitting each mediated model with a block-deranged pupil as mediator. For every derangement π ∈ *D* of block indices, pupil traces were exchanged across blocks, such that *P*^π(*b*)^ was paired with *y*^(*b*)^ and 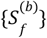, the model was re-fit, and the resulting linguistic TRFs were averaged across derangements before fold-averaging, squaring, and parcel-averaging within ROI:

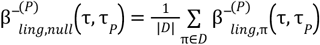

where 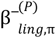 is the fold-averaged linguistic TRF from the mediated model fit with the π-deranged pupil trace. This isolates pupil-coupling from variance that would also be explained by an unrelated but pupil-like signal, such as spurious associations driven by shared temporal autocorrelation.

To equalise between-subject heterogeneity in baseline coupling magnitude, each subject’s effect surface was normalised by the root-mean-square of the squared null TRF over the tested grid (τ ≥ 0, τ_*P*_ ≥ 0),

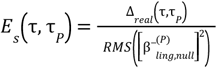

This yielded a unit-free, per-subject mass-normalised pupil-coupling effect.

#### Inference on the 2D surfaces

A two-dimensional cluster-based permutation test ^75^ was performed on *E*_*s*_ (τ, τ_*p*_) within each ROI, using sign-flip permutations (10,000 iterations) with a cluster-forming threshold of *p* = 0. 01 (one-tailed against Δ_*real*_ > 0; mne.stats.permutation_cluster_1samp_test). The stricter-than-conventional cluster-forming threshold was set following ^75^ recommendation for the relatively large search space implied by the 2D-surface × five-ROI family. To correct for multiple comparisons across the five ROIs, we constructed a family-wise max-cluster null distribution: in each of 10,000 permutations, the same subject-wise sign flips were applied to all ROIs simultaneously, within-ROI cluster masses were recomputed, and the maximum cluster mass across ROIs was retained. Observed cluster masses were referred to this single null distribution to obtain family-wise corrected p-values, *p*_*familywise*_. Within-ROI cluster p-values are reported as descriptive statistics; primary inferential decisions are based on *p*_*familywise*_.

### Statistical inference for encoding models

#### Feature importance

For cortical effects, group-level Δρ maps were tested against zero across the 360 Glasser parcels ^60^ using threshold-free cluster enhancement (TFCE; ^76^) with start = 0 and step = 0.2, embedded in a one-sample sign-flip permutation framework over Glasser-parcel adjacency (10,000 permutations, family-wise α = 0. 05; mne.stats.permutation_cluster_1samp_test with a TFCE threshold dictionary). Cortical adjacency was derived from the *fsaverage* source space and the HCP-MMP1.0 parcel-to-vertex mapping by treating two parcels as adjacent if their vertex sets shared any neighbouring vertices in the source-space adjacency graph. For cortical ROI parcels, volumetric parcels (brainstem, cerebellum) and pupillary responses (pupil diameter, pupil derivative), Δρ values were tested against zero across participants using one-sample t-tests (one-sided, against zero) where Shapiro-Wilk tests indicated approximate normality, and Wilcoxon signed-rank tests (one-sided) otherwise. Gated - Graded contrasts were tested two-sided. Effect sizes are reported as Cohen’s d (*d*) for parametric tests or the rank-biserial correlation (*r*_*rb*_) for non-parametric tests; tests were run in pingouin ^77^.

#### TRF time-courses

Cortical TRF time courses (Figure 2) summarise the response across the set of parcels that reached significance in the corresponding feature-importance TFCE map (see Statistical inference for encoding models). To collapse this multi-parcel set into a single representative time course per subject while avoiding cancellation between parcels whose TRFs share a temporal profile but differ in polarity - for example because of residual source-orientation ambiguity - we applied a sign alignment based on the singular value decomposition (SVD) of the group-mean response. The real-data TRF weights were averaged across subjects to form a parcel × lag matrix, and its first left singular vector was taken as the dominant spatial mode; the sign of each parcel’s loading on that mode defined a per-parcel polarity. These polarities were applied to every subject’s real and block-deranged TRFs - using the identical set of signs for both, so that the real-versus-null contrast remained unbiased - and the sign-aligned weights were then averaged across parcels to give one time course per subject. A single global polarity was finally fixed per response so that the dominant deflection of the group-mean time course was positive. The bilateral cerebellum response, which comprised the left and right cerebellar parcels, was collapsed by the same SVD-based sign alignment. Single-parcel responses (brainstem, pupil diameter, pupil derivative) required no alignment and were carried forward directly; the same global-polarity convention was applied to all MEG responses.

Group-level TRF time courses of features showing a significant main effect (i.e. feature importance) were tested against the block-deranged null using one-sample cluster-based permutation tests (^75^; mne.stats.permutation_cluster_1samp_test) over the (0, τ_*max*_] window, excluding pre-zero lags from inference but retaining them for display. Tests used 10,000 permutations. The cluster-forming threshold was set to *p* = 0. 05 (two-tailed for real-vs-null contrasts; tail = 0). Gated - Graded TRF contrasts were tested analogously, with sign-flip permutation on the difference-in-differences 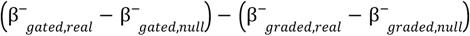 under the null of no effect (two-tailed). Significance windows are reported with their cluster p-values and peak t-statistics.

## Data and code availability

All analysis code and data to reproduce results and figures is available at ^78^. The MEG and Eye-tracking data analysed here were collected as part of a broader investigation of selective speech processing that included additional listening conditions not relevant to the present hypotheses; for full details we refer the reader to ^24^.

## Acknowledgements

We thank Manfred Seifter for his support in conducting the MEG measurements. This work was supported by a Leverhulme Project Grant (RPG-2022-358) and European Research Council (ERC) consolidator grant (101001592) under the European Union’s Horizon 2020 research and innovation programme, both awarded to C.P. Data were collected with support from the Austrian Science Fund (FWF; Doctoral College “Imaging the Mind”; W 1233-B).

## Author contributions

Q.G. designed the experiment, collected and analysed the data, generated the figures, and wrote the manuscript. A.K. contributed to data analyses and edited the manuscript. J.S. designed the experiment, collected the data, and edited the manuscript. N.W. collected the data, contributed to data analyses and edited the manuscript. C.P. acquired the funding, supervised the project, and edited the manuscript.

