## Supplemental information for "The dynamic relationship between pupil dilation and neural surprise in natural language comprehension"

### Pupil residualisation

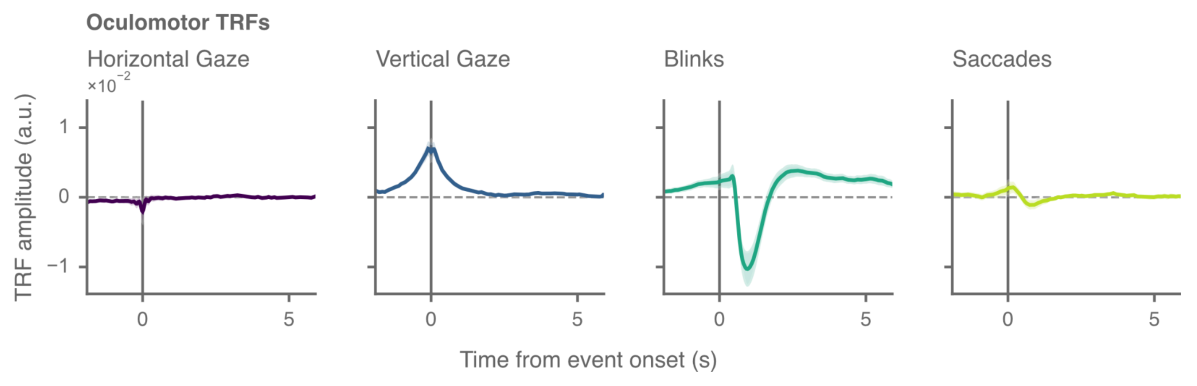


**Figure S1. Oculomotor TRFs from pupil residualisation**
Group-average TRFs from the pre-processing step in which gaze position, blink onsets, and saccade onsets were regressed out of the pupil signal using ridge regression [1] (window: −2 to +6 s; see Methods). Blink- and saccade-evoked TRFs reproduced the biphasic morphology reported by [2], with a brief constriction followed by a slower dilation after Blinks, confirming that the oculomotor regression captured the expected confounding variance. The residualised pupil signal (used in all main-text analyses) is the original signal minus this predicted oculomotor contribution. Ribbons show group mean ± SEM. N = 28.

### Cortical regions of interest

For the ROI-resolved encoding, feature-importance, and pupil-coupling analyses, we defined four a-priori left-hemisphere cortical ROIs spanning the language-processing hierarchy. The cortical ROIs were constructed as fixed groupings of HCP-MMP1.0 parcels [3] selected on anatomical and theoretical grounds to capture successive processing stages - from sensory speech input, through temporal lexical-semantic processing and temporoparietal semantic integration, to frontal prediction and update-related cortex. ROI membership was specified before analysis and was independent of the whole-brain feature-importance maps.

The early auditory ROI comprised left primary, belt, and parabelt auditory parcels, providing a sensory speech-input stage. The temporal-semantic ROI comprised left superior-temporal, superior-temporal-sulcus, and lateral-temporal parcels associated with lexical-semantic and contextual language processing. The parietal semantic-integration ROI comprised left temporo-parieto-occipital-junction (TPOJ), perisylvian-language (PSL), and inferior-parietal parcels capturing temporoparietal semantic-integration territory. The frontal ROI comprised a contiguous left inferior-frontal-to-dorsolateral-prefrontal mass spanning Broca's region (areas 44, 45) and ventrolateral prefrontal cortex (47l, p47r), the inferior frontal sulcus (IFSa, IFSp) and junction (IFJa, IFJp), and posterior middle-frontal/dorsolateral-prefrontal cortex (p9-46v, a9-46v, 46, 8C); this region covers canonical language-prediction cortex (Hagoort, 2005; Friederici, 2011) and adjacent multiple-demand regions implicated in prediction-violation processing (Brass et al., 2005; Henderson et al., 2016; Assem et al., 2020). All cortical ROIs were left-lateralised, in line with the left-dominant language network and the dominance of the left hemisphere in encoding lexical surprise and semantic prediction error in [4]. The full parcel composition of each ROI is given in Table S1.

| **Region of interest** | **n** | **HCP-MMP1.0 parcels (left hemisphere)** |
|---|---|---|
| Early auditory | 5 | A1, LBelt, MBelt, PBelt, RI |
| Temporal-semantic | 17 | A4, A5, STGa, STSdp, STSda, STSvp, STSva, TA2, TGd, TGv, TE1a, TE1p, TE1m, TE2a, TE2p, PHT, TF |
| Parietal semantic-integration | 15 | TPOJ1, TPOJ2, TPOJ3, STV, PSL, PGp, PGs, PGi, PFm, PF, PFt, PFop, IP0, IP1, IP2 |
| Frontal | 12 | 44, 45, 47l, p47r, IFSa, IFSp, IFJa, IFJp, p9-46v, a9-46v, 46, 8C |

**Table S1. Constituent HCP-MMP1.0 parcels for each cortical region of interest**
All parcels are from the left hemisphere (HCP-MMP1.0 / Glasser et al., 2016; the L_ prefix and _ROI suffix are omitted for brevity).
